# Heterogeneity of white-matter organization in the human brain

**DOI:** 10.64898/2026.03.31.714863

**Authors:** Emily Turschak, Wan-Qing Yu, Kevin Takasaki, Steven J Cook, Russel Torres, Olga Gliko, Ayana Hellevik, Kareena Villalobos, Elizabeth Guadarrama, Soumya Chatterjee, Eric Perlman, Connor Laughland, Adam Glaser, Uygar Sümbül, R Clay Reid

**Affiliations:** Allen Institute for Brain Science; Yikes, LLC; Allen Institute for Neural Dynamics

## Abstract

White-matter, which contains the long-range axons connecting different brain regions, makes up nearly half the human brain, yet the three-dimensional organization of individual axons has not been characterized. While advances in diffusion MRI have enabled macroscopic mapping of major WM pathways^1^, these methods are unable to resolve individual axons: their trajectories, density, and relative orientations^2^. Here, we present a histological and imaging pipeline optimized for post-mortem human white-matter that shows the 3D trajectories of densely stained large (diameter greater than ~1 μm) projection axons. Applying our approach to multiple centimeter-scale WM regions in an adult human brain, we observed striking regional diversity in axonal organization. Specifically, we identified distinct architectural motifs ranging from loosely packed, multi-orientation meshworks (in most superficial white matter), to laminar lattice-like structures (near the basal ganglia), to tightly packed bundles of fibers (e.g. in the corpus callosum). We speculate that these patterns reflect local adaptations to spatial constraints, axonal density, and the diversity of axonal sources and targets, offering region-specific solutions to anatomical optimization problems. These findings offer new insights into the principles shaping brain connectivity and underscore the need for regionally detailed atlases of human white matter..

---

White-matter (WM) accounts for nearly half of the human brain’s volume, a proportion that increases with brain size, making up only 12% of mouse and 25% of macaque brains^3^. While studies of neuroanatomy, primarily in experimental animals and to a lesser extent in humans, have yielded remarkable insights into mammalian brain architecture, comprehensive and high-resolution datasets characterizing WM organization in the human brain do not exist. This lack is largely due to persistent technical limitations in both microscopy and histological processing.

In fact, prior to recent advances^4,5^, axons in human WM could be traced over only very short (<1 cm) distances^6^, owing to technical limitations in both microscopy and histological processing. Direct anatomical studies of WM microstructure have long faced a tradeoff between resolution and spatial extent. For example, electron microscopy (EM) provides nanometer-level detail but is limited to small tissue volumes (≤1 mm^3^), whereas magnetic resonance imaging (MRI) can image whole brains but lacks the resolution to resolve individual axons. While large-scale initiatives such as the Human Connectome Project have generated valuable maps of macroscopic WM tracts using diffusion MRI (dMRI) and tractography^1,7^, these methods yield only indirect, model-based estimates of axon trajectories^8,9^.

These technical barriers have long been recognized as a fundamental obstacle to understanding the human brain. Over 30 years ago, Francis Crick and Edward Jones^10^ decried “The backwardness of human neuroanatomy” compared to pathway tracing studies in experimental animals. Their blunt assessment still rings true today: *“The shameful answer is that we do not have such detailed maps because, for obvious reasons, most of the experimental methods used on the macaque brain cannot be used on humans*.*”* After decades of research, our understanding of the microstructure of human WM remains extremely limited, as we still lack techniques capable of tracing long-range axons at single-fiber resolution or revealing their fine-scale spatial organization within human WM.

Advances in histological processing (including hydrogel-based tissue expansion^11–17^ and optical clearing^18–23^ and in microscopy techniques such as high-throughput lightsheet imaging^24^ have begun to overcome these barriers^4^. However, human WM remains particularly challenging due to its dense architecture and light-scattering properties. Continued development of scalable, single-axon resolution imaging pipelines are essential for building anatomically accurate, regionally detailed maps of human WM organization, an endeavor that stands to significantly advance our understanding of brain connectivity in health and disease^25,26^.

We ultimately aim to construct a projection map of the human brain at single-axon resolution. As a step toward this goal, we demonstrate the diverse local organization of axon trajectories across human WM regions, extending observations from previous histological studies and inferences from dMRI^27–32^. Our results demonstrate that human WM is far from uniform in its microarchitecture, but instead is organized into strikingly distinct regional motifs that reflect local anatomical and functional demands. These findings establish a foundation for understanding lower-resolution approaches, such as dMRI, and for mapping brain connectivity at single-axon resolution.

## Methods

We developed a pipeline combining antibody labeling, optical clearing, hydrogel expansion, and light-sheet imaging to resolve individual axons across centimeter-scale volumes of post-mortem human WM (see Supplementary Methods). The tissue processing protocol integrated elements from tissue preservation strategies (eg. SHIELD^23^, LICONN^16^), tissue delipidation and optical clearing (eg., LifeCanvas delipidation: Park 2019; CUBIC^22^), and hydrogel-based isotropic expansion (eg. ExM^11^, ELAST^14^, LICONN^16^. Flash-frozen coronal slabs (0.5 cm) of an adult human brain^33^ were fixed and cut into ~0.6×0.6×0.5 cm tissue volumes (“punchouts”) (Fig. 1A), and subsequently sectioned at 500 μm (Fig. 1B) on a Leica VT 1000S vibratome. Sections were incubated in a polyepoxide-based stabilization solution (SHIELD) and delipidated with Clear+ buffer (*LifeCanvas Technologies*^*23,34*^. Neurofilament heavy protein (NFH) was labeled using our adapted immunolabeling protocol. NFH was selected based on its established use for myelinated long-range projection axons^35–38^. Control experiments with viral labeling of excitatory neurons in macaques demonstrated that all virally labeled cortical axons were also labeled with NF antibodies (data not shown *[can be included in revision])* Samples were then embedded in a tissue-hydrogel matrix and isotropically expanded to 3x their original size (Fig. 1C, D). Expanded gels were adhered to polylysine-coated glass slides and mounted to a sample arm using epoxy-protected magnets, followed by imaging with an ExA-SPIM^24^, a high-speed, large-field-of-view light-sheet microscope.

**Figure 1.**
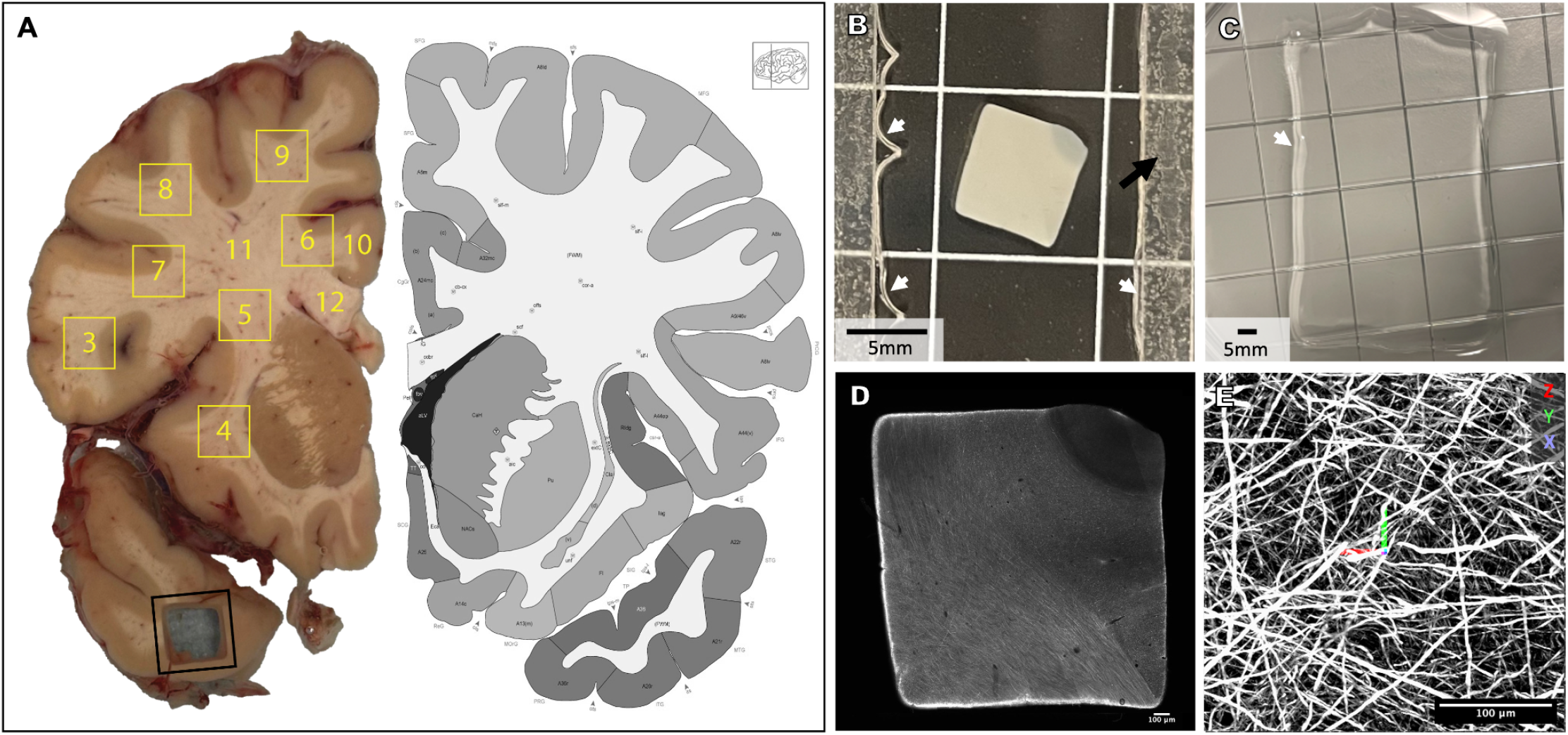
(A) Fixed hemi-coronal brain slab (~0.5 cm thick), with a representative ~0.6 cm^2^ punchout removed from the temporal lobe (black outline) to illustrate the sampling approach. Yellow outlines and labels indicate the locations of the sampled regions. Punchout 2 is not shown. Punchouts 3-9 were obtained using the punch tool, whereas 11 and 12 were dissected from the remaining tissue with scalpel cuts. Corresponding atlas image from Allen Institute website [resource](https://atlas.brain-map.org/atlas?atlas=265297126#atlas=265297126&plate=111257849&structure=10328&x=45088.52237654321&y=42718.311149691355&zoom=-7&resolution=153.97&z=3). (B) 500 μm section from punchout #6. White arrows = gel border. Black arrow = spacer for polymerization chamber. (C) Cleared tissue-hydrogel matrix, expanded in water. White arrow = gel border. (D) Confocal overview of polymerized section, pre-expansion. (E) ExA-SPIM lightsheet image example. Scale bars relative to actual size, not corrected for expansion.

## Results

We examined NFH labeled axons across samples to identify shared features and characterize regional and structural variability in axonal organization. NFH labeling produced strong signal in axons, with minimal background staining. There was considerable spacing between axons, which enabled most with larger caliber (above ~1.5 μm in diameter) to be readily traced manually with few ambiguities. Here, we address local WM structure rather than long-range connectivity, so detailed analysis of axon segmentation and traceability is deferred to a more technical report^5^.

Our findings highlight striking regional diversity in axonal organization across human WM, which we refer to here as multi-orientation, laminar/orthogonal, and bundled (Fig. 2A-C). The statistics of local axonal organization can be quantified with structure tensor analysis, a method widely applied in image processing, computer vision, and biomedical imaging, to characterize fiber orientations^39–41^. Structure tensor analysis enables color-coding of axons based on their local orientation, with red coding for medio-lateral, green for dorso-ventral, and blue for antero-posterior orientations (Fig. 2D-F). The distribution of orientations at a location yields a 3-dimensional orientation distribution function (ODF; Fig. 2G-I). In the case of the multi-orientation domains (Fig. 2A, D) the ODF typically has multiple broad peaks (Fig. 2G), while the laminar/orthogonal domains (Fig. 2B, E) has two peaks (Fig. 2H).

**Figure 2.**
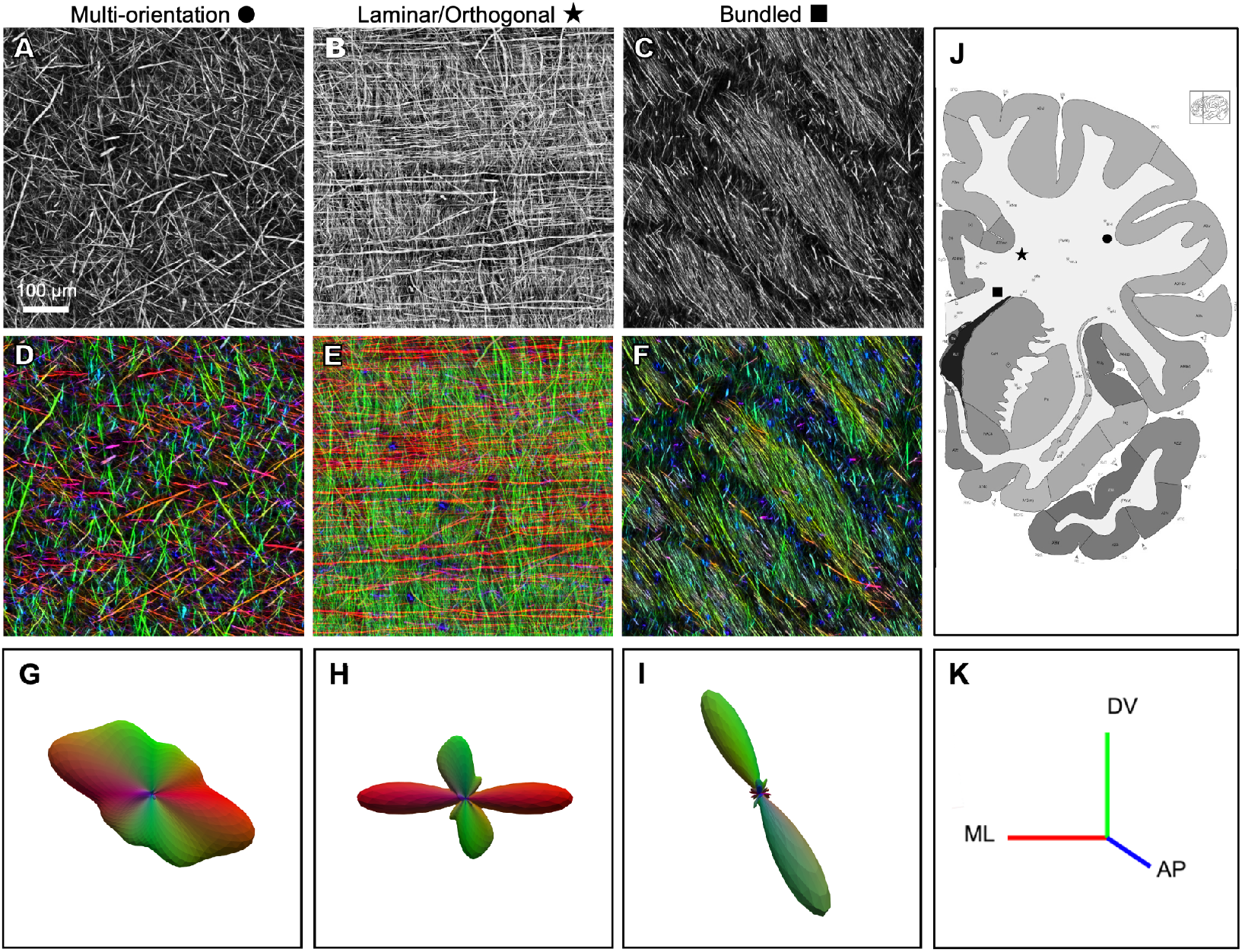
Visualizing the data with structure tensor analysis (A-C) Representative examples of three distinct organizational motifs: (A) multi-orientation, (B) laminar/orthogonal, and (C) bundled, shown in grayscale. (D-F) Corresponding orientation maps derived from structure tensor analysis, with color indicating local axonal orientation (D: multi-orientation, E: laminar/orthogonal, F: bundled). (G-I) ODFs for each example, illustrating the distribution of axonal orientations (G: multi-orientation, H: laminar/orthogonal, I: bundled). (J) Coronal atlas section corresponding approximately to the sampled tissue slab. (K) Structure tensor color key, with red indicating medio-lateral, green dorso-ventral, and blue antero-posterior orientations.

The multi-orientation organization was found in superficial WM, near the cortical surface, characterized by relatively low density of NFH-labeled axons (Fig. 2A), multiple orientations (Fig. 2D), and an absence of discernible local bundling (Fig. 2J), resulting in a seemingly random three-dimensional meshwork. Further from the cortical surface, such as the broad expanse of WM known as the centrum semiovale, the multi-orientation mesh-like structure was observed, although with mixtures of orientations that varied gradually across the region (see below).

In other WM regions, we observed several varieties of more structured axonal organization. The laminar/orthogonal organization was seen at the border of the centrum semiovale, near the basal ganglia and corpus callosum. Most axons had one of two orthogonal orientations^42^, with a laminar organization (which predominates in mouse WM^43^) that appeared almost woven. (Fig. 2B, J, and see below). This may represent an alternate organizational solution for routing axons through constrained spaces, possibly balancing structural coherence with the need to accommodate multiple trajectory angles within the same volume.

The bundled organization was seen in several locations. Near the basal ganglia, (Fig. 2C, F) we observed intertwined bundles, each with largely parallel axons, but with slightly different orientations and different apparent densities. One hypothesis is that the sparser bundles contain a mixture of axons with differential antibody staining. The bundled organization was also seen in the corpus callosum, where all axons had similar orientations, but organized in smaller bundles of nearly parallel fibers (see Fig. 3G, below). This organization is consistent with prior descriptions of the corpus callosum as a high-throughput commissural pathway, composed primarily of long-range axons that connect homologous cortical areas across hemispheres^27,32,44–48^.

**Figure 3.**
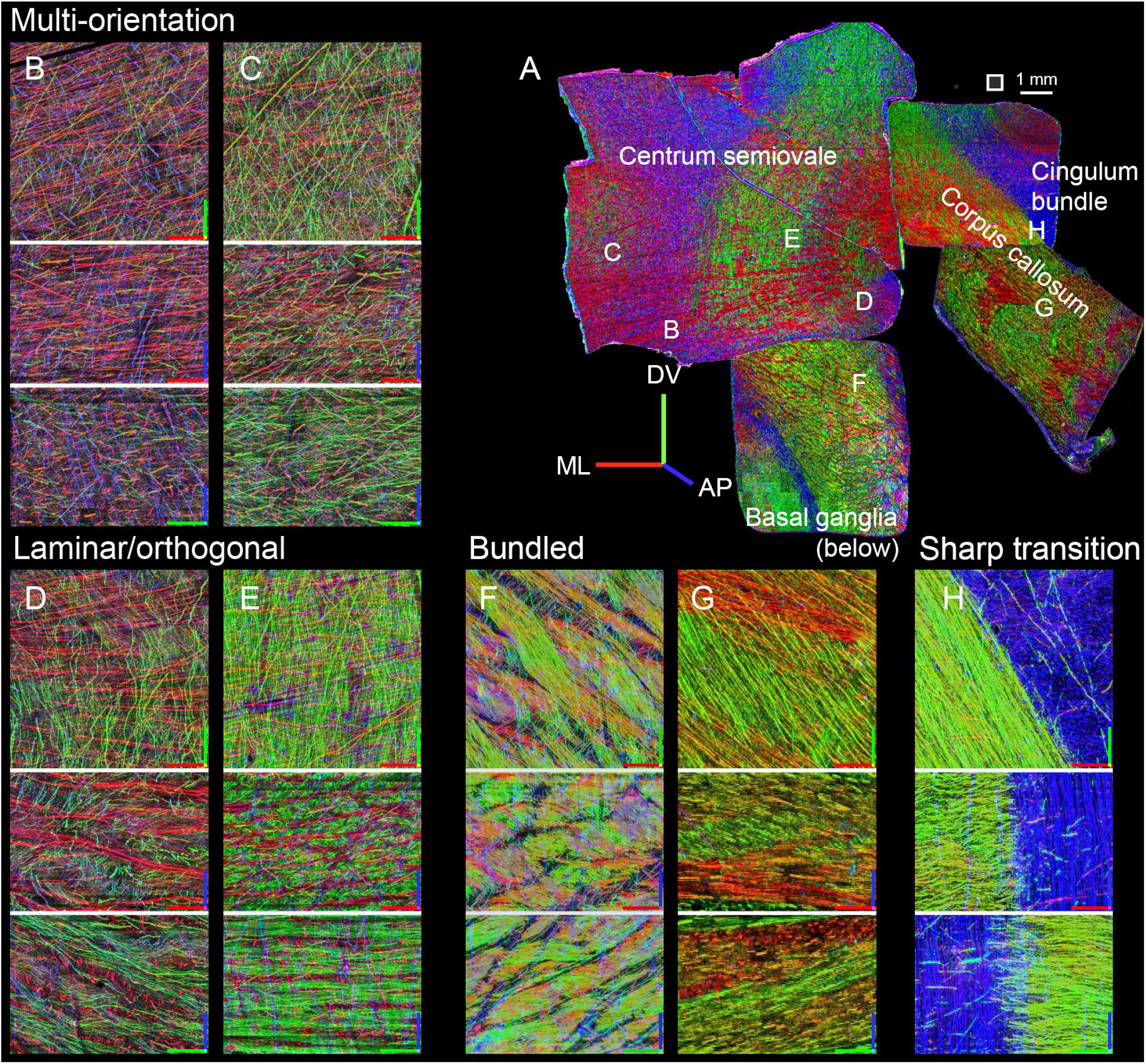
Examples of each type of WM organization. (A) Montage of the four central punchouts(11, 6, 12, and 5), manually arranged to approximate their spatial positions within the original tissue slab.(Fig. 1A). Square next to scale bar indicates the size of regions (500 µm) in the remaining panels . (B-H) Three orthogonal views through the middle of 500×500×350 µm image stack, shown as maximum intensities, projected through 60 µm (B-E) or 30 µm (F-H; because of the higher density of bundled axons). Axes are color-coded as follows: red=medio-lateral, green=dorso-ventral, blue=antero-posterior. The red-green plane represents the top-down view of the samples. Axis length = 100. Panels are grouped by configuration. (B-C): Multi-orientation, with individual, ungrouped axons coursing in different directions. ; (D, E): Laminar/orthogonal, with lamination visible in the bottom two projections, in the AP/blue direction. (F, G): Bundled, above the basal ganglia and in corpus callosum, respectively. (H) Border between bundled organization in the corpus callosum and multi-orientation (but with a predominance of the antero-posterior orientation) in cingulum bundle.

To further illustrate these different modes of organization, we offer examples from the four most central punchouts, including the corpus callosum, its border with the cingulum bundle, the centrum semiovale, and its inferior bordering region near the basal ganglia (punchouts 12, 6, 11, and 5, respectively). The multi-orientation meshwork was the predominant pattern observed in portions of the centrum semiovale near the cortical surface (Fig. 3B, C), while the laminar/orthogonal pattern was seen in adjacent positions nearer the corpus callosum (Fig. 3D, E). As noted above, bundled organization was seen near the basal ganglia, just superior to the external capsule (Fig. 3F) and within the corpus callosum (Fig. 3G). Axon density increased progressively from multi-orientation to laminar/orthogonal to bundled configurations. At the interface between the corpus callosum and adjacent WM near the cingulum bundle, we observed an abrupt transition between the tightly bundled axonal architecture of the corpus callosum and the cingulum bundle (Fig. 3H), which despite the high density of antero-posterior axons, had isolated axons coursing with other orientations.

To quantify these observations, we analyzed the same four most central punchouts (Fig. 4A) by first calculating the structure tensor orientation distributions in 0.4 × 0.4 mm regions (Fig. 4A). We quantified these distributions using generalized fractional anisotropy (GFA), a metric commonly used in dMRI that reflects the degree of axonal alignment, ranging from 0 (all orientations equally represented) to 1 (a single dominant orientation).

**Figure 4.**
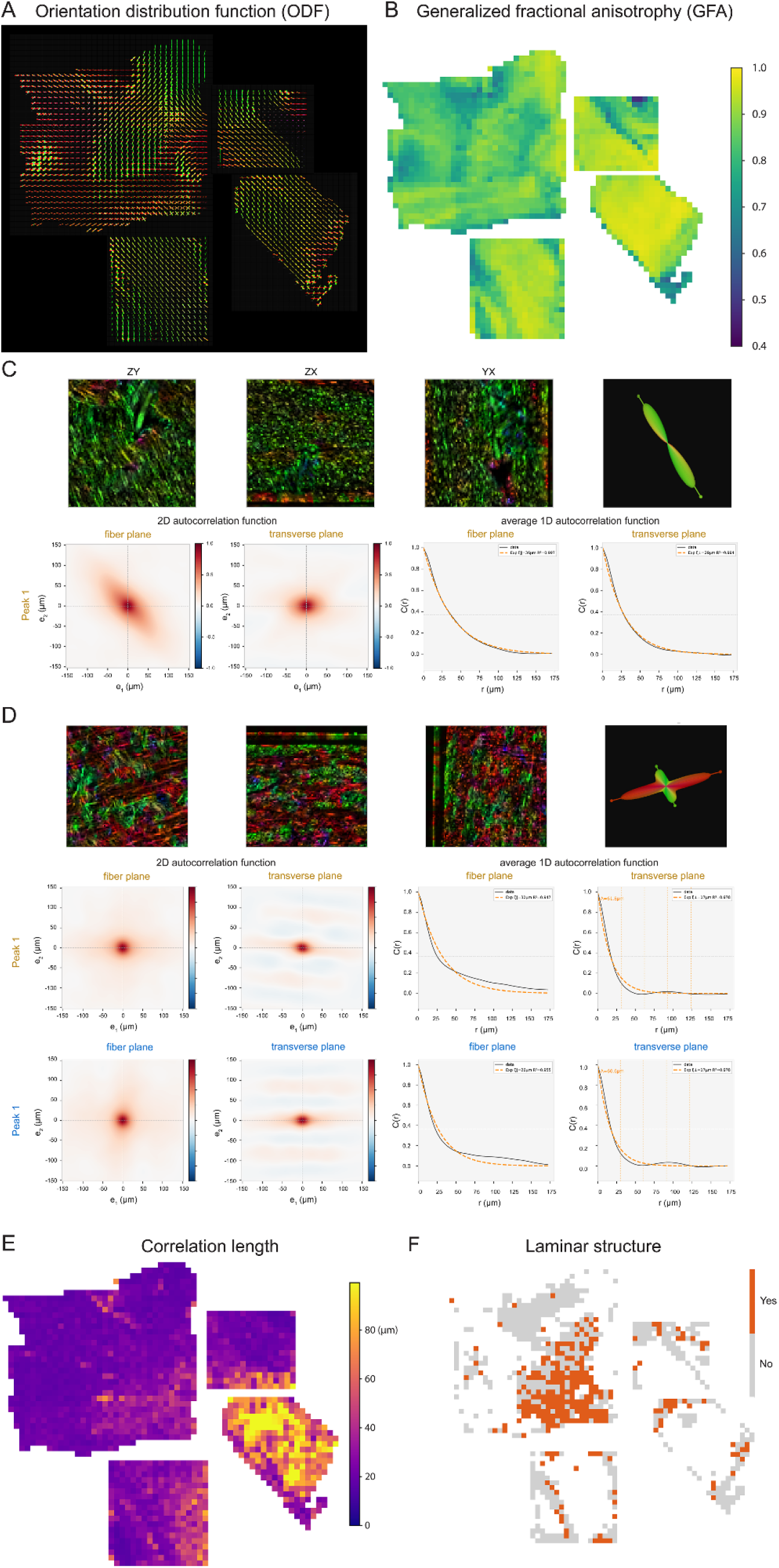
Multiscale quantitative characterization of fiber organization across human WM regions. (A) 3D orientation distribution functions (ODF) glyphs per 0.5 mm^3^ block. Glyph shape reflects the degree of directional coherence, with elongated forms indicating aligned fiber populations and multi-lobed forms indicating crossing or dispersed fiber arrangements. (B) Spatial heatmap of generalized fractional anisotropy (GFA; range 0.4-1.0) quantifying regional variations in orientation coherence. Higher values (yellow) indicate strongly aligned fiber tracts; lower values (purple) reflect isotropic or complex fiber architecture. (C, D) Characterization of local spatial arrangements of fibers via autocorrelation function analysis. (C) Representative coherent tract motif with tightly aligned fibers, yielding a single dominant ODF peak and smooth exponential decay in both 2D and radially averaged 1D autocorrelations in the fiber and transverse planes. (D) A representative crossing orthogonal fiber with laminar structure, yielding a two-peak ODF. Notably, both the 2D and average 1D autocorrelation in the transverse plane exhibit spatial oscillations, providing quantitative evidence of a periodic, laminar-like fiber arrangement rather than random intermingling. (E) Spatial heatmap of maximum autocorrelation length, revealing the physical scale over which the fiber orientation remains coherent. Regions with long correlation lengths (yellow) correspond to large, organized fiber bundles; short correlation lengths (purple) reflect rapid change of spatial organization. (F) Binary map showing the multi-peak regions containing laminar structure (orange = present, grey = absent)

Mapping this metric across these punchouts (Fig. 4B) revealed distinct spatial domains, distinguishing the highly parallel arrangement (high GFA) in the corpus callosum from regions of complex, multidirectional dispersion (low GFA). To quantitatively determine the spatial organization of fibers across regions, we applied autocorrelation functions on vector fields derived from structure tensor analysis (Fig. 4C-D). In bundled regions characterized by a single dominant ODF peak, such as the corpus callosum, the autocorrelation exhibited a monotonic decay without periodicity (Fig. 4C). Conversely, in laminar/orthogonal regions containing alternating layers of orthogonally oriented fibers, the 2D and average 1D autocorrelation in the transverse plane revealed distinct spatial oscillations (Fig. 4D). Extending this analysis across the four punchouts, we generated spatial heatmaps of the maximum autocorrelation length. This metric mathematically defines the spatial scale of organization, ranging from approximately 20 µm to over 90 µm. Finally, to delineate the spatial extent of these laminar domains, we generated a binary map based on the presence or absence of periodic autocorrelation oscillations, requiring detection of at least two orientation peaks in the ODF. This analysis clearly revealed that the region of the centrum semiovale nearest the callosum and the basal ganglia (lower right) comprises widespread, contiguous domains of structured, laminar-like fibers across several millimeters of tissue.

## Discussion

We applied our pipeline to centimeter-scale WM regions from an adult human brain. Our findings highlight striking regional diversity in axonal organization across the WM, including a multi-orientation meshwork in superficial WM, near the cerebral cortex (Fig. 2A), orthogonally oriented trajectories arranged in a laminar organization trajectories in regions near the basal ganglia (Fig. 2B), and tightly bundled, coherent fiber arrangements in and near the corpus callosum (Fig. 2C).

Histological studies have long suggested that WM microstructure varies across brain regions^27,28,43,49,50^, reflecting underlying circuit specialization. For example, differences in axon diameter, packing density, and myelin sheath thickness are well-documented in the human corpus callosum ^27,32,45,47^, which is well known for having densely bundled axons with a limited range of orientations^44,48^. Similar diversity in axon caliber has also been observed across some cortical and commissural pathways^46^, aligning with the notion that WM architecture is shaped by the distinct functional demands of each tract.

While the biological basis for these organizational motifs remains incompletely understood, several hypotheses grounded in anatomical and geometric principles may help explain this variation. One likely factor is the spatial density and physical constraints of the local WM environment. In regions of high axonal density, bundled, parallel fiber arrangements may offer an efficient solution to packing large numbers of axons into compact volumes. Prior theoretical work suggests that such bundling may reduce wiring length and metabolic cost while maintaining large diameters required for high conduction velocity^28,51,52^. This is consistent with observations in the corpus callosum and internal capsule, where tightly aligned fibers likely reflect a drive toward wiring optimization under strong spatial constraints.

In other regions (such as in portions of the centrum semiovale, Figs. 2A, 3C, 3D), we observed a laminar organization: stacks of layers with alternating, nearly orthogonal fiber orientations. Rather than forming parallel fascicles, which by definition are not planar structures, the layers maintained relatively consistent spacing and orientations (as quantified in Fig. 4), although the layers frequently intermingled (Fig. 3C, D), producing woven or lattice-like configurations. These layers present an alternate organizational solution for routing axons through moderately dense regions, possibly balancing structural coherence with the need to accommodate multiple angles within the same volume. The predominance of near-orthogonal trajectories was noted previously^30,42^, but the laminar organization appears novel.

Finally, in areas with lower axonal density (such as subcortical WM) axons may be less tightly constrained to achieve dense packing. This can give rise to more heterogeneous, multi-orientation configurations, where axons traverse the tissue along independently varying paths. This uncorrelated meshwork organization may present an optimal solution for accommodating multiple pathways that overlap en route to diverse cortical or subcortical targets^46^. Such arrangements resemble the intersection zones described in dMRI studies as sites of high orientation dispersion^53^. Theoretical models have proposed that axonal trajectories reflect a balance between minimizing total wiring length and preserving topographic specificity, which has been termed the “wiring economy” principle^45,54–56^. From this perspective, the observed regional differences in organization may emerge from local solutions to a high-dimensional spatial optimization problem, constrained by both physical space and the complexity of circuit topology.

Although further work is needed to formalize and test these hypotheses, our findings demonstrate that human WM is highly variable in its organization, exhibiting region-specific microarchitectural adaptations that may reflect differences in functional demands, developmental trajectories, and spatial constraints. These patterns will have important implications in multiple areas: (1) for understanding how large-scale brain networks are physically constructed, (2) for interpreting disruptions to WM organization in neurodevelopmental and neurodegenerative disease^25,26^, (3) to inform the calibration and interpretation of lower-resolution techniques, such as dMRI and optical coherence tomography, and (4) to guide neurosurgical approaches that depend on accurate delineation of white-matter architecture.

## SUPPLEMENTARY METHODS

### Histology

#### Tissue Source, Slab Preparation, and Sectioning

Postmortem human tissue from an adult male donor (61 years old, Hispanic, with no known neuropsychiatric or neurological conditions) was obtained from the San Diego Medical Examiner’s Office. The brain was processed by bisecting at the midline, as described in^33^, and coronal slabs approximately 0.5 cm thick were embedded in alginate, rapidly frozen in a dry ice-isopentane slurry, vacuum sealed, and stored at -80°C. When retrieved for processing, the frozen slab was drop-fixed in ice-cold 4% paraformaldehyde (PFA) and 10% acrylamide in phosphate-buffered saline (PBS) for 24 hours at 4°C with gentle agitation (as adapted from the LICONN protocol). The fixed tissue was then rinsed overnight in 100mM glycine in 1×PBS at 4°C to quench residual aldehydes^34^, followed by 3 rinses (>15min each) in 1×PBS at 4 °C with gentle agitation.

Individual 1cm^3^ regions of interest (Fig. 1A) were sampled from the slab using a square punch tool. Each tissue block was embedded in 4% agarose in 1X PBS and sectioned at 500 μm on a Leica VT 1000S vibratome for downstream processing. Individual sections were transferred to a 15 mL tube, where all subsequent processing steps were carried out to minimize tissue handling and reduce contamination risk. Sections remained in their respective tubes until polymerization.

#### Post-Fixation

*(adapted from Park et al., 2018)*

Tissue sections were incubated in a polyepoxide-based stabilization method adapted from the SHIELD protocol^23^, which preserves protein antigenicity and tissue architecture during downstream clearing and expansion. Sections were incubated in SHIELD-Off buffer for 72 hours at 4°C with gentle agitation, followed by incubation in SHIELD-On (*LifeCanvas Technologies*) buffer for 24 hours at 37°C with gentle agitation. This step minimizes tissue degradation and molecular loss during subsequent processing, enabling consistent preservation across large volumes of human WM. After SHIELD stabilization, sections were rinsed in 1xPBS three times for 15 minutes each.

#### Delipidation

Tissue sections were delipidated using Clear+ buffer (*LifeCanvas Technologies*^*23,34*^) for 2 weeks at 37 °C with gentle agitation. Buffer volume was supplemented after one week to ensure adequate penetration (3mL initial + 3mL additional). Following delipidation, samples were rinsed in 1×PBS containing 0.02% sodium azide to remove residual clearing agents. Sodium azide was included in this rinse step to prevent microbial growth.

#### Antibody labeling

Blocking was performed in NGSTU-azide buffer (prepared by diluting 10×PBS to 1× with NGSTU-azide buffer) for 24 hours at room temperature. Urea was included in the blocking and primary antibody buffer solutions to enhance tissue permeability, an approach inspired by CUBIC protocols^33^. Primary antibody incubation was carried out using anti-NFH antibody (see below) (1:500) in NGSTU-azide buffer. Samples were incubated sequentially under the following conditions: 2mL for 1 week at 4°C, an additional 2mL for 1 week at room temperature, and a final addition of 2mL for 1 week at 37°C, all with gentle shaking. Following primary antibody incubation, tissue was rinsed with 5mL of 1×PBS-azide a minimum of four times for 15 minutes each. For the final rinse, tissue was incubated in 10mL of 1×PBS-azide overnight at 4°C. Secondary antibody incubation used 1:250 goat anti-mouse Alexa Fluor 488 antibody, see below) in NGST-azide buffer. Urea was not included in the secondary antibody buffer solution due to concerns of fluorescence quenching. Sections were incubated with 4mL of secondary antibody solution for 1 week at room temperature, with an additional 2 mL added at the one-week mark, followed by incubation for 1 more week at 37°C (final volume: 6 mL). Following secondary antibody incubation, tissue was rinsed with 5 mL of 1×PBS a minimum of five times for 15 minutes each, then transferred to 10 mL of 1×PBS overnight at 4°C. Sodium azide was removed from this rinse step to prevent interference with downstream hydrogel polymerization (it is highly reactive, which could potentially interfere with or destabilize the desired reaction products). All incubations steps, including rinses, were done with gentle shaking.

#### Chemical Anchoring

*(adapted from Tillberg et al., 2016)*

Acryloyl-X, SE (AcX) was used to covalently anchor amine-containing biomolecules to link them to the expanding polymer matrix (see below). AcX was first resuspended in anhydrous DMSO to a concentration of 10 mg/mL, aliquoted, and stored at −20 °C in a desiccated environment for up to 2 months.

For anchoring, tissue sections were incubated in 0.1 mg/mL AcX diluted in 1xPBS, pH6, for at least 72 hours at 4°C with gentle agitation. AcX treatment introduces acrylamide moieties to endogenous proteins, enabling subsequent covalent linkage to the polymer network during hydrogel polymerization. Tissue was then rinsed in 1xPBS five times for 15 minutes each.

#### Expanding Hydrogel Monomer Incubation

Prior to gelation, the expanding hydrogel monomer stock solution^16^ was thawed on ice. For each 1 mL of monomer solution, 15 μL of VA-044^14^, a thermal initiator, was added.

The solution was vortexed to ensure homogeneity, and 1 mL was added to each tissue section (1 cm × 1 cm × 0.5 mm) in their respective 15 mL conical tubes, ensuring full coverage of the tissue. Samples were incubated for at least 72 hours at 4 °C with gentle agitation to allow thorough diffusion of the monomer into the tissue.

Immediately prior to polymerization, tube caps were removed and samples were placed in a vacuum desiccation chamber for 15 minutes to degas the solution and minimize oxygen inhibition of polymerization.

#### Chamber construction and polymerization

Polymerization chambers were constructed by placing a 0.5mm adhesive spacer (16×26mm inner; 22×30mm outer; see Fig. 1B) on a microscope slide. The chamber was filled with monomer solution and tissue was placed in the center of the spacer. A 24×55mm #1 coverslip was then carefully placed on top of the spacer, ensuring that no bubbles were trapped in the solution. Chambers were placed in a 10cm petri dish, which was subsequently gently purged with nitrogen gas for 30 seconds. The perimeter of the petri dishes was parafilmed, and dishes were placed in a 37°C incubator for two hours without agitation. Once polymerized, each tissue section was imaged on a confocal microscope while still in the chamber to obtain an overview image prior to removal (see Fig. 1D). This post-polymerization, pre-expansion overview image served as a reference for precise size measurements before enzymatic digestion and tissue expansion, enabling accurate calculation of the final expansion factor.

#### Enzymatic digestion

Tissue-hydrogel matrices were removed from polymerization chambers and transferred to 60×15mm dishes with 10mL 1:100 Proteinase K in the digestion buffer. Samples were then incubated at 37°C overnight with gentle shaking.

#### Expansion

After enzymatic digestion, gels were carefully transferred to 150×150×25mm dishes and rinsed in 1xPBS three times for 15 minutes each. Gels were transferred into excess volumes of Milli-Q water and incubated for 15 minutes to 2 hours to initiate expansion. This water exchange step was repeated 3-5 times using fresh Milli-Q water until the sample reached ~4x its original size. Gels were then equilibrated in 0.5x PBS, resulting in a 3x final expansion for lightsheet imaging.

## Microscopy

### Sample mounting and large-format lightsheet microscopy

A custom expansion-assisted selective plane illumination (ExA-SPIM) lightsheet microscope was constructed according to the design and resources described in the original publication ^24^ and modified for imaging of 500 μm-thick, 3x expanded sections immersed in 0.5x PBS. Expanded gels were adhered to glass slides coated with polylysine which were then mounted with epoxy-protected magnets to a sample arm tilted at 60 degrees and actuated along the tilted axis with an LS-50 motorized linear stage (Applied Scientific Instrumentation). The tilted stage enabled constant velocity scanning through the lightsheet for tile acquisition and the three-axis motorized stage system (MS-8000 and FTP-100, Applied Scientific Instrumentation) effected gross repositioning of the section between tiles. Microscope hardware was controlled by a custom lightsheet device control and image acquisition software package (Voxel, Allen Institute for Neural Dynamics) and image data was compressed and written directly to the next-generation file format zarr version 3 by an open-source data writer (acquire-zarr, Chan-Zuckerberg Institute). Zarr datasets were concurrently uploaded during acquisition to an on-premises, enterprise all-flash data platform (VAST) with acquisition metadata for downstream processing with the HPC cluster.

Individual physical voxels were ~0.75 μm, nearly isotropic, which corresponds to ~0.25 μm relative to the unexpanded tissue, given the 3x expansion.

## Computation

### Image data preprocessing: deskewing and downsampling for multiresolution pyramid

Raw data volumes generated by the virtual-V ExASPIM acquisition were skewed with respect to the tissue coordinates and raw images are composed entirely of full resolution voxels. Data preprocessing on the HPC cluster resolves these shortcomings by generating output zarr datasets that have been deskewed into the usual XYZ tissue coordinates. The zarr files contained a downsampled image pyramid (MIP levels) which facilitates visualizing and processing large volumes of data. MIP0 voxels are ~0.25 μm, relative to the original tissue, MIP1 voxels are ~0.50 μm, etc.

### Section preprocessing: stitching

Deskewed tiles were first assembled into a volume in the unified coordinate space of the section by applying translation offsets according to the predefined position of the stage in the acquisition. These offsets were further refined by registering image data in the overlapping regions of the tiles. A custom stitching algorithm uses a combination of SIFT and template matching to define informative points from highly downsampled data around which sub-voxel phase correlation is applied to the highest resolution to extract empirical spatial offsets beyond those defined by the coordinates of the microscope stage^5^. These offsets were recorded as metadata allowing the deskewed tiles to be viewed and manipulated as a single, “stitched volume” by standard big data tools, such as Neuroglancer (https://github.com/google/neuroglancer), as well as the downstream processing pipeline.

### MIP4: fusion/transformation to get anatomical orientation

For fusion of adjacent tiles, we designed a module suited for the translation of multiple adjacent tiles. Each tile is placed into a common coordinate space using precomputed translation offsets, whether stage or registration derived, which specify the position of each tile’s origin within the global volume, and written into the appropriate location in a single output volume. The output is a single consolidated volume ready for downstream processing steps.

There were several punchout volumes adjacent to one another, but processed without regard to their original orientation. The impact of this on the structure tensor analysis is a mismatch of orientation values for axon traces between adjacent volumes. To resolve this issue, we rotated those volumes using a rotation degree derived from visual inspection in Neuroglancer.

### Data analysis

For quantitative analysis of fiber architecture, MIP4 image datasets were divided into 100×100×100 voxel cubes. We utilized structure tensor analysis^57^ to extract primary fiber orientations from local image intensity gradients, which were then projected onto a 6,500-point spherical space. This generated discrete orientation distribution functions for each cube, allowing us to calculate generalized fractional anisotropy and identify orientation peaks separated at least by 45°. To evaluate the spatial arrangement of these fiber tracts, we calculated the 3D autocorrelation function from the local orientation tensor field. The spatial scale of structural coherence, correlation length, was derived by fitting axial autocorrelation profiles to an exponential decay model. Finally, to systematically map regions of laminar structure of cross fibers without relying on visual inspection, we screened transverse ACF profiles for spatial periodicity. This classification required strict spectral evidence, including ≥1.5 full cycles in the spatial domain coupled with robust signal-to-noise thresholds across both 1D and 2D fourier domains. Tissue cubes with low fractional anisotropy and at the border of the tissues were omitted from all spatial maps.

## Neuroglancer links

### Raw

#### PO12

https://neuroglancer-demo.appspot.com/#! https://apex-connects.s3.us-east-2.amazonaws.com/axonal_connectomics/contrast_adjusted/H17_PO12_S4_20250501_V2/state2.json

#### PO6

https://neuroglancer-demo.appspot.com/#! https://apex-connects.s3.us-east-2.amazonaws.com/axonal_connectomics/contrast_adjusted/H17_PO6_S3_20250422/state2.json

#### PO5

https://neuroglancer-demo.appspot.com/#! https://apex-connects.s3.us-east-2.amazonaws.com/axonal_connectomics/contrast_adjusted/H17_PO5_S8_20250410/state2.json

#### PO11

https://neuroglancer-demo.appspot.com/#! https://apex-connects.s3.us-east-2.amazonaws.com/axonal_connectomics/contrast_adjusted/H17_PO11_S8_20250408/state.json

### Structure tensor MIP4

#### PO12

https://neuroglancer-demo.appspot.com/#! https://apex-connects.s3.us-east-2.amazonaws.com/axonal_connectomics/contrast_adjusted/H17_PO12_S4_20250501/equalized_fused/state.json

#### PO6

https://neuroglancer-demo.appspot.com/#! https://apex-connects.s3.us-east-2.amazonaws.com/axonal_connectomics/contrast_adjusted/H17_PO6_S3_20250422/equalized_fused/state.json

#### PO5

https://neuroglancer-demo.appspot.com/#! https://apex-connects.s3.us-east-2.amazonaws.com/axonal_connectomics/contrast_adjusted/H17_PO5_S8_20250410/equalized_fused/state.json

#### PO11

https://neuroglancer-demo.appspot.com/#! https://apex-connects.s3.us-east-2.amazonaws.com/axonal_connectomics/contrast_adjusted/H17_PO11_S8_20250408/equalized_fused_st/state2.json

### Fused (lower res)

https://neuroglancer-demo.appspot.com/#! https://apex-connects.s3.us-east-2.amazonaws.com/axonal_connectomics/contrast_adjusted/smlori11-Big-mv960-160_rot100/equalized_fused/state.json

## Supplemental: Materials and equipment

### Reagents

- Paraformaldehyde 32% aqueous solution EM grade (Electron Microscopy Sciences, #15714-5)
- Glycine (Sigma-Aldrich, #G7126)
- PBS 10x buffer pH7.4 1L (Life Technologies, #AM9625)
- SHIELD kit (SHIELD Epoxy, Buffer, and ON solutions), (LifeCanvas Technologies, SH-250)
- Clear+ delipidation buffer (LifeCanvas Technologies, #DB)
- Agarose (VWR, #IB70042)
- Normal Goat Serum (NGS), (Vector Labs # S-1000)
- Triton X-100 (Sigma-Aldrich, # X100)
- Urea (Sigma-Aldrich, #U2709)
- Sodium Azide, 5% (Fisher Scientific, #71448-16)
- Rabbit anti-neurofilament heavy chain (NF200 or NFH) antibody (Sigma-Aldrich, N4142)
- Goat Anti-Rabbit IgG H&L (Alexa Fluor® 488), (Abcam, ab15007)
- DMSO, Anhydrous (ThermoFisher Scientific, D12345)
- Acryloyl-X (Acx), (ThermoFisher Scientific, A20770)
- Acrylamide (Sigma-Aldrich, #A9099)
- Sodium acrylate, (AK Scientific, #7446-81-3)
- N,N′-(1,2-Dihydroxyethylene) bisacrylamide (Sigma-Aldrich, #294381)
- VA-044 (Fisher Scientific, #NC0632395)
- Sodium dodecyl sulfate (SDS), (Sigma-Aldrich, #L4509)
- 1M Tris-HCI, pH 8.0 (ThermoFisher Scientific, #15568025)
- Proteinase K (ThermoFisher Scientific, #EO0491)
- Poly-L-Lysine (Sigma-Aldrich, #P4707)

### Materials/equipment

- ~1 × 1 cm hollow square punch tool (Yizzvb, https://www.amazon.com/dp/B08T94ZLXM)
- Vibratome (Leica VT 1000S)
- Corning® 15ml conical tubes (Millipore Sigma, #CLS430055)
- 50 mL conical tubes
- Epredia™ Richard-Allan Scientific™ Cover Glass, #1, 24×55 (Fisher Scientific,124405)
- iSpacer 0.5mm, (SUNJin Lab Co., #IS002)
- Corning® microscope slides (Millipore Sigma, #CLS294875X25)
- 150X25mm dishes (Fisher Scientific, 08-772-6)
- 100mm×15mm (Fisher Scientific, #263991)
- 60×15mm dishes (Carolina, #741246)
- Olympus FLUOVIEW FV3000
- Lightsheet (ExASPIM)

### Solutions

## Blocking Buffer and Primary Antibody Buffer

The blocking and primary antibody buffer (NGSTU-azide) was prepared in 1×PBS with the following composition:

- 5% (v/v) normal goat serum
- 0.6% (v/v) Triton X-100
- 4M urea
- 0.02% (v/v) sodium azide

All components were mixed thoroughly until fully dissolved. Buffer was stored at 4°C for up to 1 month.

## Secondary Antibody Buffer

The secondary antibody buffer (NGSTU-azide) was prepared in 1×PBS with the following composition:

- 5% (v/v) normal goat serum
- 0.6% (v/v) Triton X-100
- 0.02% (v/v) sodium azide

## Expanding Hydrogel Monomer Solution

The expanding hydrogel monomer stock solution was prepared using the following components *(adapted from Tavakoli et al., 2025):*

- 10% (w/v) acrylamide
- 12.5% (w/v) sodium acrylate
- 0.075% (w/v) N,N′-bisacrylamide

All components were dissolved in Milli-Q water. The solution was vortexed thoroughly and centrifuged at 4,500 × g for 5 minutes to remove particulate matter. The resulting supernatant was transferred to a fresh 50 mL conical tube, aliquoted into 1 mL volumes, and stored at −20 °C for up to one month.

## Digestion Buffer

*(adapted from Tillberg et al., 2016)*

- 100 mM Tris base
- 5% (v/v) Triton X-100
- 1% (w/v) sodium dodecyl sulfate (SDS)

All components were dissolved in Milli-Q water, and the solution was mixed thoroughly until fully solubilized. The buffer was prepared fresh or stored at room temperature for short-term use. Prior to use, the buffer was equilibrated to 37 °C. This solution facilitates enzymatic digestion and mechanical disruption of the tissue-hydrogel composite to allow uniform and isotropic expansion.

